# CosinorTest: An R Shiny App for Cosinor-model-based circadian and differential analysis in transcriptomic applications

**DOI:** 10.1101/2024.11.06.622324

**Authors:** Haocheng Ding, Yutao Zhang, Lingsong Meng, Chengguo Xing, Karyn A. Esser, Zhiguang Huo

## Abstract

**Summary:** ‘CosinorTest’ application is an interactive R Shiny tool designed to facilitate circadian and differential circadian analysis with transcriptomic data using Cosinor-model-based methods. This novel application integrates multiple major statistical algorithms to identify circadian rhythmicity in gene expression data and enables the comparison of differential circadian patterns between two experimental conditions. Key features of the ‘CosinorTest’ app include circadian rhythmicity detection, differential patterns assessment, circadian and differential analyses with repeated measurement, and interactive data visualization, all contributing to a comprehensive understanding of underlying biological mechanisms. As the application doing all comprehensive circadian analysis using the Cosinor model, the ‘CosinorTest’ app fills a crucial gap in the field of circadian biology and transcriptomics, providing a powerful and user-friendly platform for researchers, especially those without profound programming skills to explore the circadian gene expression regulation, and further advance circadian research.

**Availability and implementation:** CosinorTest is freely available at https://circadiananalysis.shinyapps.io/circadianapp/.

**Supplementary information:** Supplementary files are available at *bioRχiv* online.

## 1 Introduction

Circadian rhythms are endogenous biological processes that follow a *∼* 24 hours cycle and regulate various physiological and behavioral functions in living organisms, including humans. Circadian rhythms play a vital role in regulating sleep-wake cycles, hormone secretion, body temperature, and metabolism (Badia et al., 1991; Jung et al., 2010; Cagnacci et al., 1992; DIJK et al., 1992). These functions are crucial for maintaining internal synchronization with the external environment, ensuring optimal functioning and health. Disruption of these rhythms has been linked to various health issues, including sleep disorders, mood disorders, cardiovascular diseases, and metabolic disorders. Another important research question is the differential circadian analysis. It is the comparison of circadian rhythms between different experimental conditions. Previous epidemiology and animal studies have shown that the disruption in clock and circadian gene expression might trigger diseases including cancer (Sancar and Van Gelder, 2021; Ballesta et al., 2017), type 2 diabetes (Stenvers et al., 2019), sleep disorder (Möller-Levet et al., 2013), major depression disorder (Li et al., 2013), aging (Chen et al., 2016), schizophrenia (Seney et al., 2019), and Alzheimer’s disease (Lim et al., 2017).

As circadian omics studies have emerged as a significant field in recent years (see Figure 1 for number of publications), various methods have been developed for detecting circadian rhythmicity and differential circadian patterns. In terms of circadian rhythmicity detection, the Cosinor-based model assumes the expression level of a gene is a cosine function of the circadian time (Cornelissen, 2014). This model is biologically interpretable and provides accurate statistical inferences under Gaussian assumptions, as shown by Ding et al. (2021). Other parametric models, such as Lomb-Scargle periodograms (Glynn et al., 2006) and COSOPT (Straume, 2004), can detect oscillating genes with irregular shape by assuming mixtures of sine and cosine curves with distinct frequencies. Cosinor2 (Cornelissen, 2014) and CosinorPy (Moškon, 2020) are capable of detecting circadian rhythmicity with repeated measurements. Nonparametric models, including ARSER (Yang and Su, 2010), RAIN (Thaben and Westermark, 2014), and JTK CYCLE (Hughes et al., 2010), do not rely on any model assumptions, making them powerful tools for capturing irregular curve shapes. MetaCycle (Wu et al., 2016) aims to combine results from ARSER, JTK CYCLE and Lomb-Scargle using meta-analysis via Fisher’s method. Both parametric and nonparametric algorithms have been widely adopted in circadian transcriptomic studies, and several review studies have systematically compared the performance of these algorithms (Hughes et al., 2017; Mei et al., 2020; Laloum and Robinson-Rechavi, 2020).

**Figure 1:**
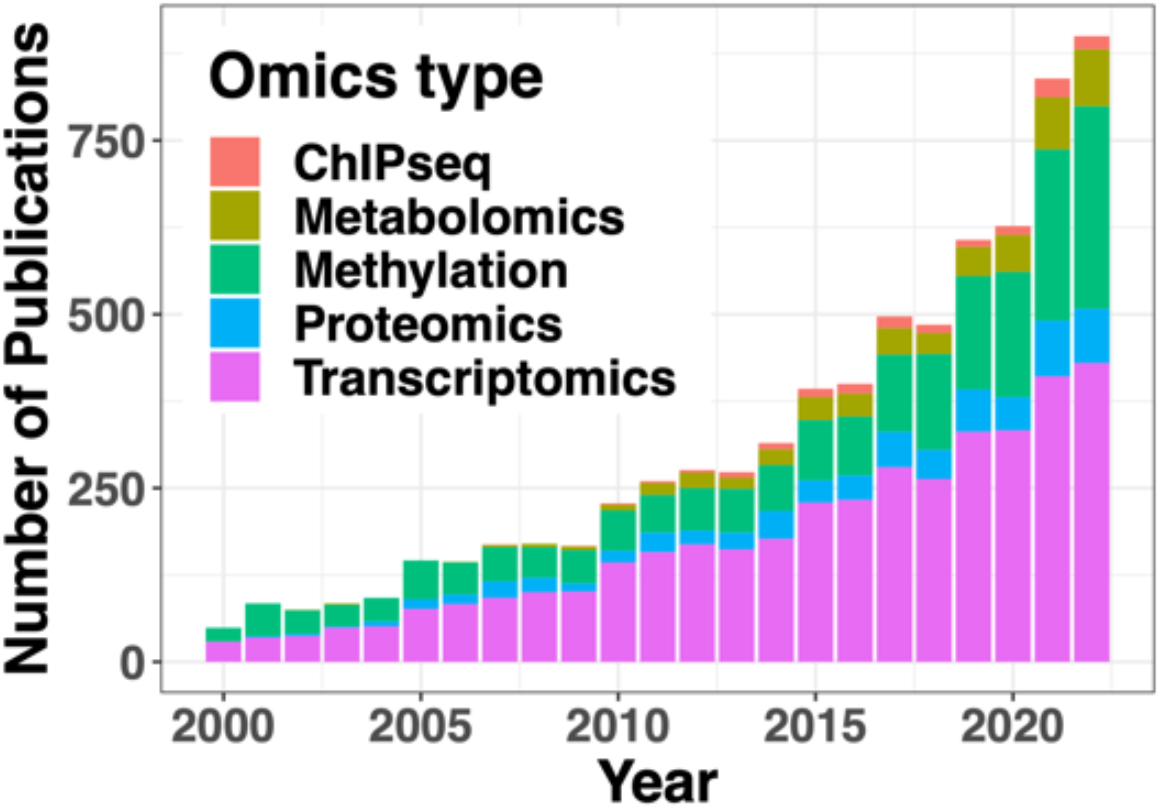
Annual number of publications on PubMed that contain the keywords “circadian/clock” and one of the following omics types: “ChIPseq”, “Metabolomics”, “Methylation”, “Proteomics”, or “Transcriptomics”.

**Figure 2:**
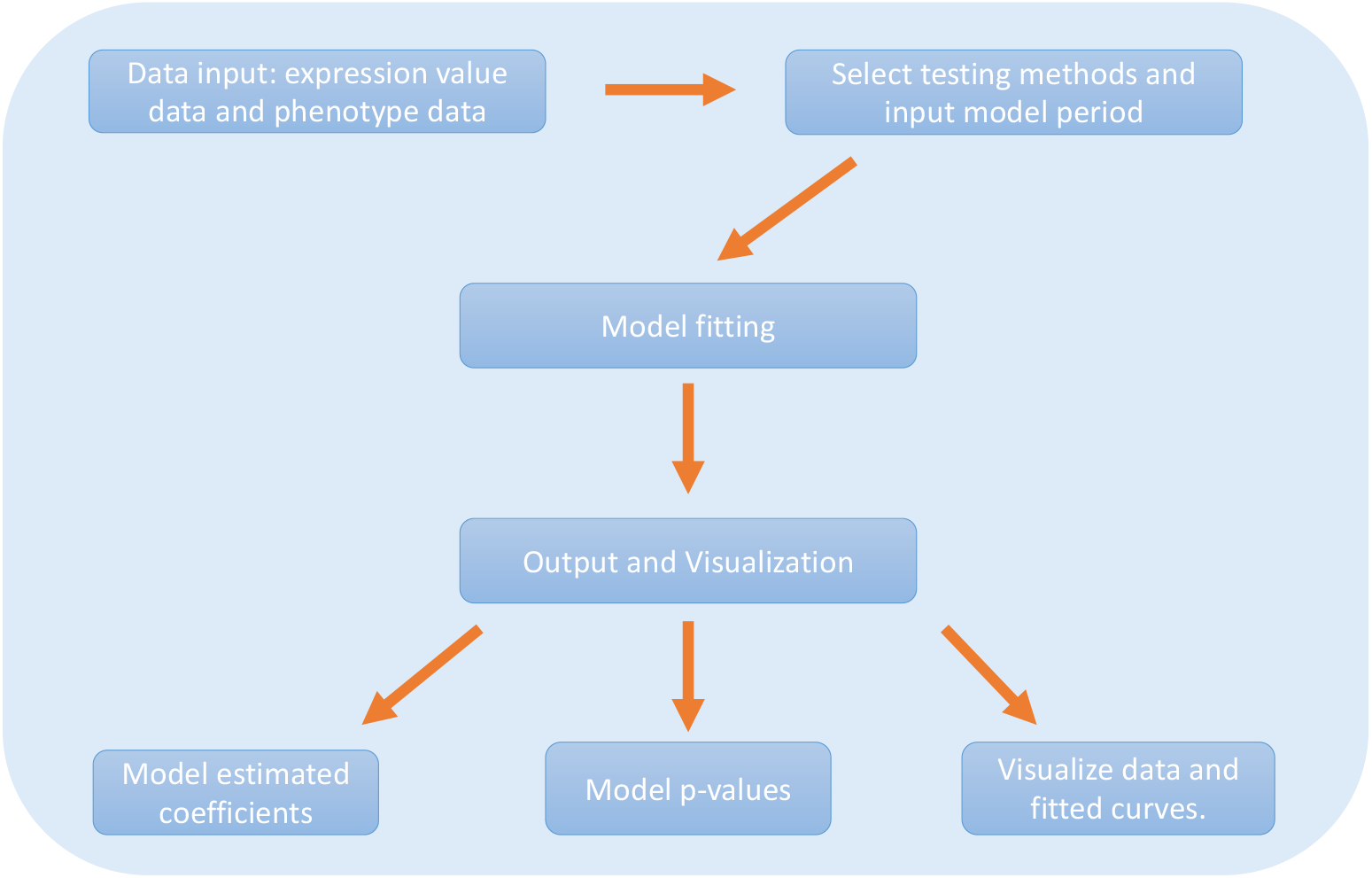
Workflow of CosinorTest.

In our R Shiny App, we mainly focused on Cosinor models because of the following reasons: (i) The biological rationale is that the circadian rhythm is amenable to adapting to the cycles in the environment (Takahashi, 2016). Since the light-dark cycle is the leading environmental factor (Scheving and Halberg, 1980), a Cosinor wave model is widely used to emulate the Cosinor cycle of the day-light intensity. (ii) The Cosinor model is parsimonious and its parameters, such as amplitude, phase (time at circadian peak), and MESOR (mid-line estimating statistic of rhythm), are biologically meaningful and widely accepted in circadian studies (Seney et al., 2019; Chen et al., 2016; Ketchesin et al., 2020). (iii) The parametric form of the Cosinor model allows it to be extended to perform differential analysis or repeated measurement analysis, and these extensions are very commonly encountered in the literature, but are not available based on non-parametric methods.

In addition, the circadian field involves identifying differential circadian patterns under different experimental conditions (Hughey and Butte, 2016; Hsu and Harmer, 2012; Möller-Levet et al., 2013). Numerous epidemiological and animal studies have established connections between disruptions in clock and circadian gene expression and various diseases, such as cancer, type II diabetes (Stenvers et al., 2019), sleep disorders (Möller-Levet et al., 2013), major depressive disorder (Li et al., 2013), aging (Chen et al., 2016), schizophrenia (Seney et al., 2019), and Alzheimer’s disease (Lim et al., 2017). In the literature, all existing methods are based on the Cosinor model, which can be categorized into two major types. The first type examines whether there is a difference in the overall circadian pattern, with the null hypothesis being that the cosine curve fittings are identical across two experimental conditions. Representative statistical methods falling under this first type include RobustDODR (Thaben and Westermark, 2016), HANOVA (Thaben and Westermark, 2016) and LimoRhyde (Singer and Hughey, 2019). On the other hand, the second type of Cosinor-based methods aims to pinpoint the exact differences, such as changes in amplitude, peak time, MESOR, or goodness of fitness. Representative statistical methods in this second category include the permutation approach (Chen et al., 2016), CircaCompare (Parsons et al., 2020), cosinoRmixedeffects (Hou et al., 2022), ASSIST (Wang et al., 2003), Cosinor2 (Cornelissen, 2014) and diffCircadian (Ding et al., 2021).

Another important research question is related to the repeatedly measured data in a circadian experiment – multiple measurement of the same subject at different circadian time. For example, Lundell et al. (2020) recruited eleven overweight or obese men to examine the impact of restricted feeding on circadian transcriptome of skeletal muscle, with each participant being repeatedly measured every 4 hours over a 24-hours cycle. This type of circadian experiment with repeated measurement is prevalent in the literature (Perrin et al., 2018; Möller-Levet et al., 2013; Blume et al., 2017; Saner et al., 2021). These repeated measurement from the same subject are naturally correlated, and failing to consider these correlation structures may result in poor statistical powers in detecting circadian rhythmicity.

Though these methods are promising, most of these algorithms were implement in R software. In the vast expanse of biological research, practitioners come from diverse backgrounds. Not every biologist or practitioner has formal training in programming or the nuances of intricate software like R. Though undoubtedly potent, the learning curve for R can be a deterrent for many. Even for those well-versed in R, a user-friendly interface can substantially reduce the time spent on data analysis. Instead of writing and debugging code, users can invest their valuable time in interpreting results and furthering their research. In light of the above points, several web-based applications have been designed for circadian rhythmicity detection, such as Nitecap (Brooks et al., 2021), DiscoRhythm (Carlucci et al., 2021), and CIRCADA (Cenek et al., 2020). While these applications boast a well-designed, user-friendly interface, some notable research gaps still exist to hinder biologists and practitioners to fully employ the existing methods, that is, none of them support differential circadian analysis or repeated measurement analysis. To address these functional application gaps, our browser-based application, CosinorTest, offers multiple testing options for both circadian and differential circadian analyses. It is freely available on the R Shiny server at ‘www.shinyapps.io’.

## 2 Program overview

Our R Shiny app provide comprehensive Cosinor-based tools to handle various scenarios that are commonly encountered in circadian analysis. Two functional modules are included to perform circadian and differential analyses, respectively. Each module has two sub-modules for the purpose of repeated measurement data, with total of four testing conditions: (i) circadian rhythmicity detection with independent samples; (ii) circadian rhythmicity detection with repeated measured samples; (iii) differential circadian analysis with independent samples and (iv) differential circadian analysis with repeated measured samples. In each testing condition, there are three output sections including: uploaded data, analysis result table and data fitting visualization. More detail and function of each module are described in the Supplementary Material.

## 3 Discussion

Our web-based Cosinor model for circadian analysis (CosinorTest) is a free web application with four testing sub-modules to perform circadian and differential circadian analyses with either independent or repeated measured data. All our methods are based on the Cosinor model because of its biological relevance, clear interpretability, and its capability to accommodate differential circadian analyses and repeated measurement data, areas to which non-parametric methods cannot be extended. It is capable of handling multiple genes’ expression value data simultaneously and can be easily used without requiring a programming background. For each testing module, there are several available methods that can be chosen, and the results can be visualized based on users’ specifications. In terms of its performance, it has been evaluated in our previous studies for developing testing methods in circadian and differential circadian detection (Ding et al. (2021), Ding et al. (2022)). At present, our application necessitates data that satisfies the assumption of normality. If the Gaussian assumptions are violated, this can lead to either an inflated or deflated type I error rate. Although our methods are based on the Cosinor model, which offers precise inference and straightforward interpretation, it is not suited for irregular model shapes. To address these concerns, we are considering the inclusion of a robust test statistic in our upcoming updates. Due to limited space, a comprehensive user tutorial is provided in the Supplementary Material.

## 5 Acknowledgement

Large Language Models (i.e., chatGPT) were used to correct written text (spell checkers, grammar checkers).

