## Supplementary Material for "CosinorTest: An R Shiny App for Cosinor-model-based circadian and differential analysis in transcriptomic applications"

Tutorial

### 1. Introduction

Cosine model for circadian analysis is an R shiny application for scientist to perform circadian rhythmicity detection and differential circadian analysis on gene expression value data. The application is publicly available at <https://circadiananalysis.shinyapps.io/circadianapp/>.

### 2. Navigation page

After launch the application, there is a navigation bar menu including 5 sections: (1) About, (2) Circadian rhythmicity detection module, (3) Differential circadian analysis module, (4) Tutorial and (5) Troubleshooting. In terms of questions and bug reporting, please contact maintainer Haocheng Ding.

### 3. Application modules

#### Module 1: Circadian rhythmicity detection

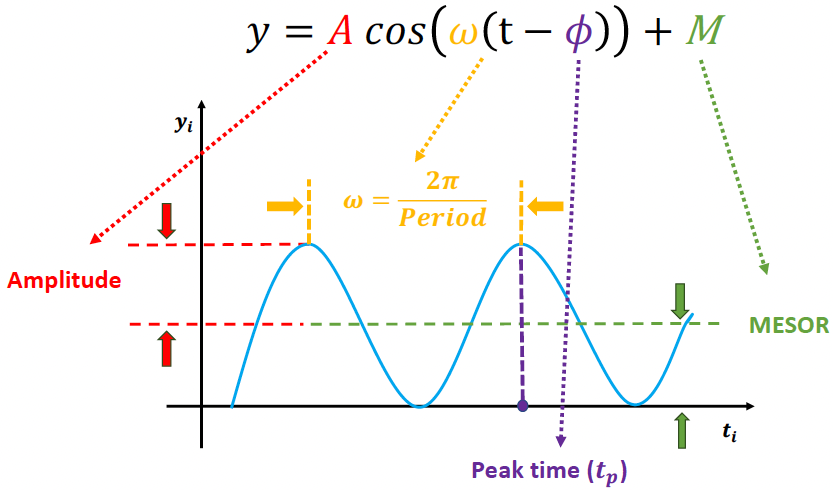

Figure 1 Cosine curve wave model for circadian rhythmicity detection

The model assumes that the relationship between the gene expression level and the circadian time fits a cosine wave curve. As shown in Figure 1, denote y as the expression value for a gene; t as the circadian time; M as the MESOR (midline estimating statistic of rhythm) level; A is the amplitude. ω is the frequency of the sinusoidal wave, where ω = 2π/*Period*. φ is the phase of the cosine wave curve.

**Hypothesis testing**

The equation can be re-written as $y_{i} = E sin(\omega t_{i}) + F cos(\omega t_{i}) + M + \varepsilon_{i}$, where E = Acos(ωφ), and F = Asin(ωφ). The hypothesis setting for testing the existence circadian rhythmicity is $H_{0} : E = F = 0 v.s. H_{A} : E\neq0 or F\neq0$.

##### a. with independent samples

**Data preparation**

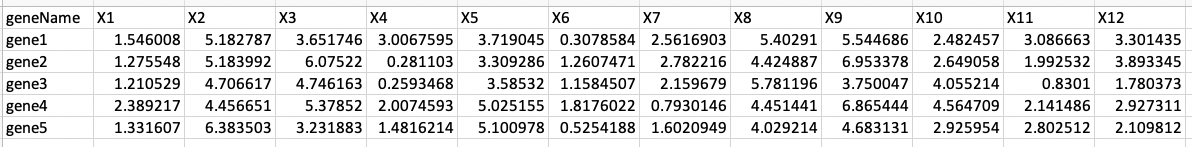

Figure 2: Sample expression data for circadian rhythmicity detection with independent samples

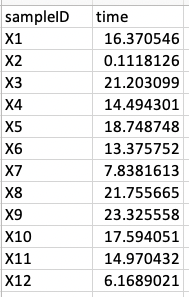

Figure 3: Sample phenotype data for circadian rhythmicity detection with independent samples

In the expression data, each row represents one gene, and each column represents one time point. The sample IDs (X1, X2, …) should match with the sample IDs in phenotype data. The unit of time variable in the phenotype data is hour(s).

**Input**

- Expression data: gene expression value data in .csv format (see Figure 1).
- Phenotype data: phenotype data in .csv format (see Figure 2).
- Period: period of gene expression value data, default is 24.

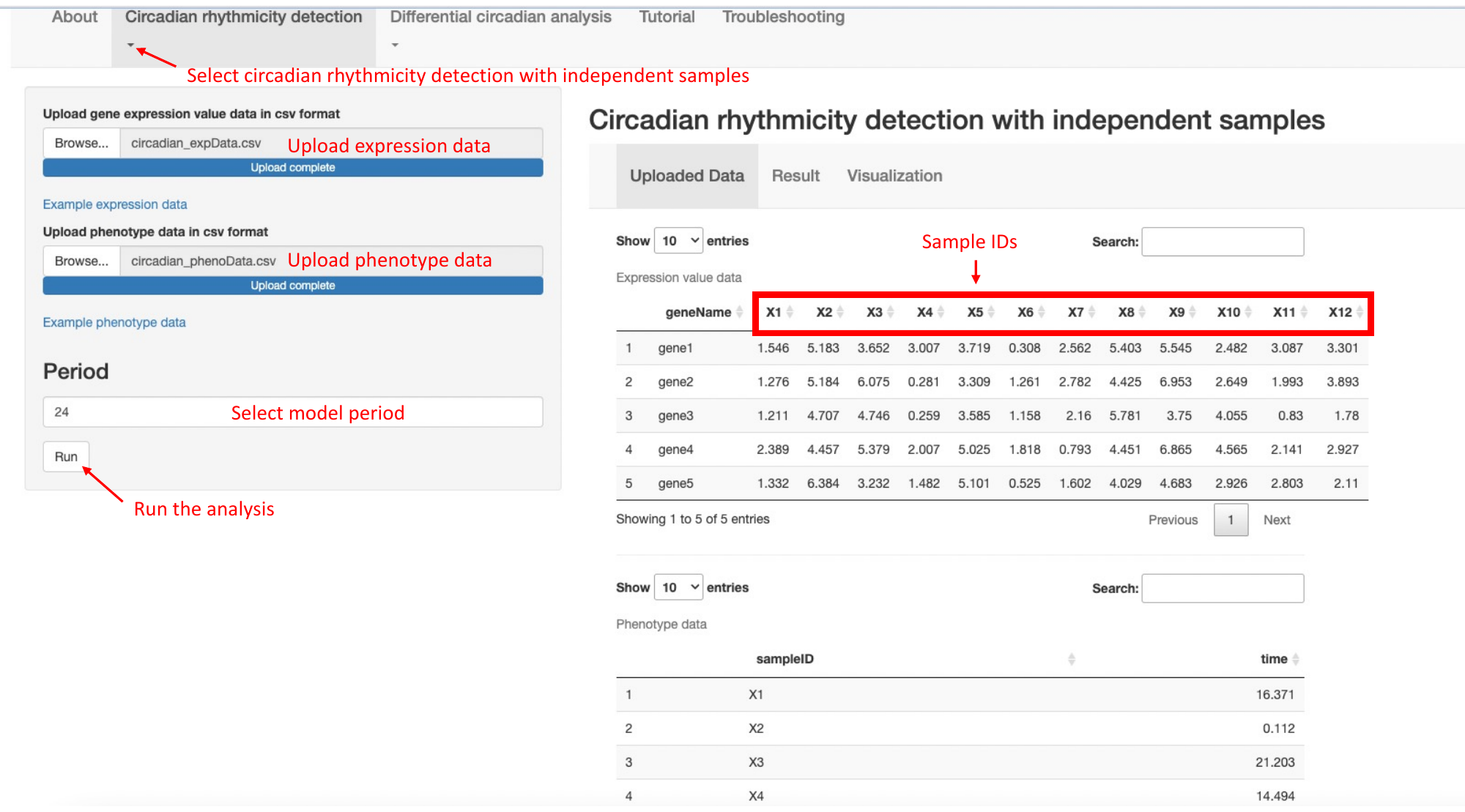

Figure 4: Circadian rhythmicity detection with independent samples after uploaded expression and phenotype data

**Run**

In left input panel, first upload both gene expression value data and phenotype data. Then input the period (default is 24) and click the ‘Run’ button (see Figure 3). Click the ‘Result’ box in the right panel, it will show the estimates summary table. The ‘Download Result’ button allows you to down the result table (see Figure 4). In addition, click ‘Visualization’ box allows user to visualize the model fitting result. To plot the result, simply input interested gene name then click the ‘Plot’ button and download the plot with ‘Download Figure’ button (see Figure 5).

**Output**

A result table including cosine model parameter estimates and statistics (see Figure 4), detailed explanations are shown below:

- geneName: gene name.
- amplitude: estimated amplitude.
- peakTime: estimated peak time of the cosine curve.
- MESOR: estimated value of MESOR level (vertical shift).
- pvalue: p-value from the test.
- R_square: Pseudo R^2^ defined as (total sum of square – residual sum of square)/total sum of square.

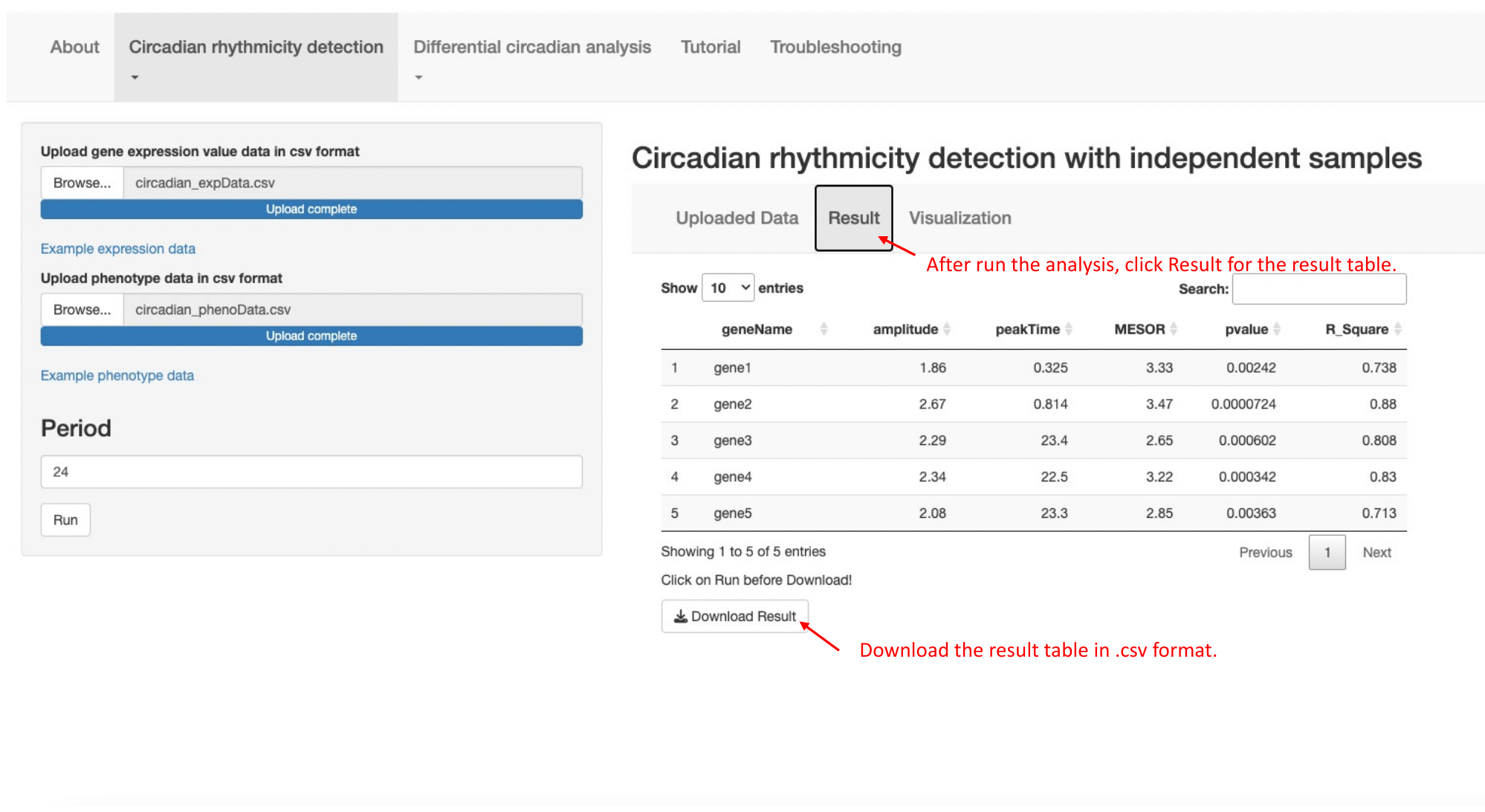

Figure 5: Result table of circadian rhythmicity detection with independent samples

Circadian gene plot: model fitting plot of interested circadian gene.

- Gene name: interested gene name.

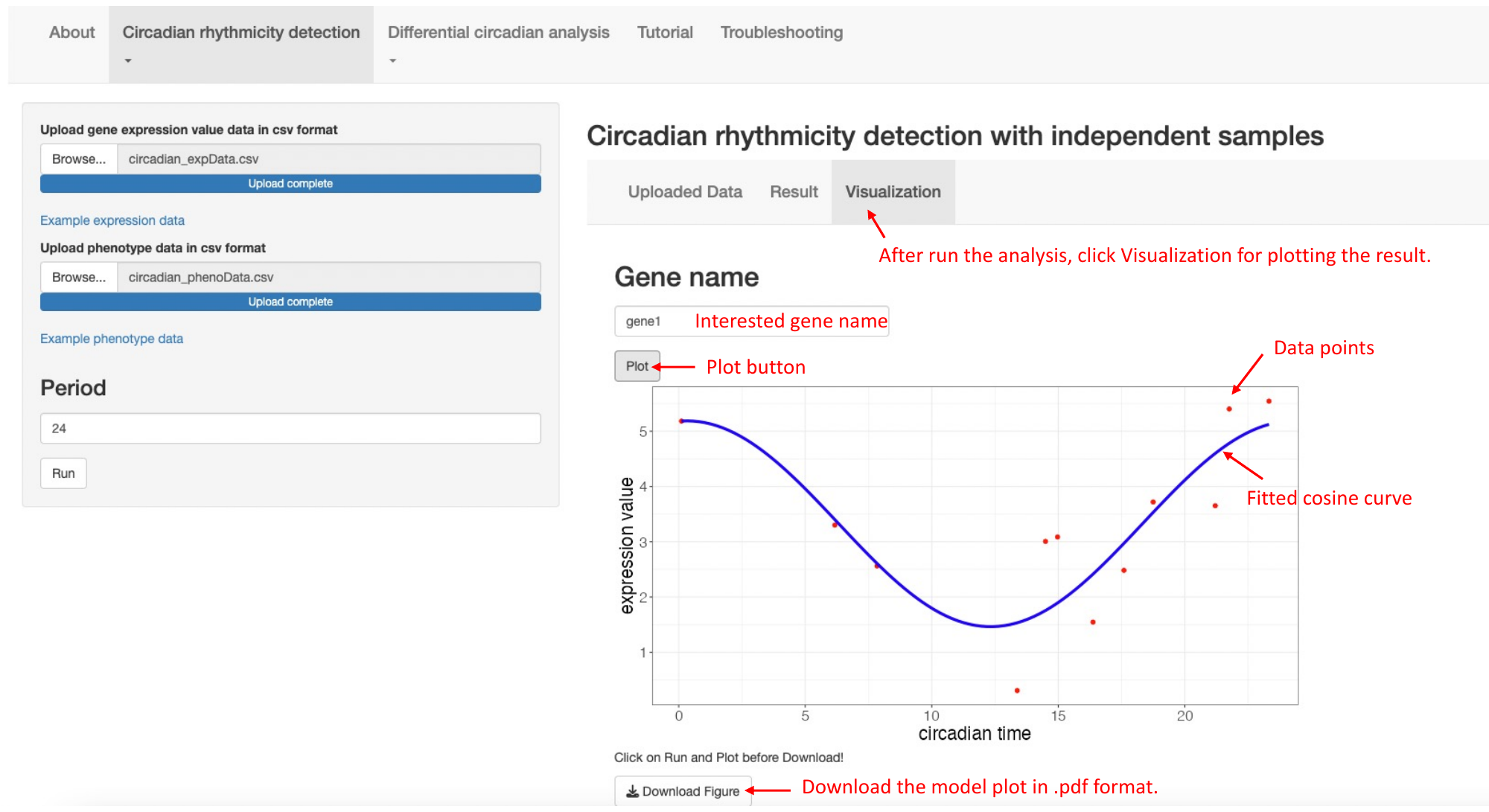

Figure 6: Visualization result of circadian rhythmicity detection with independent samples

##### b. with repeated measurement

**Data preparation**

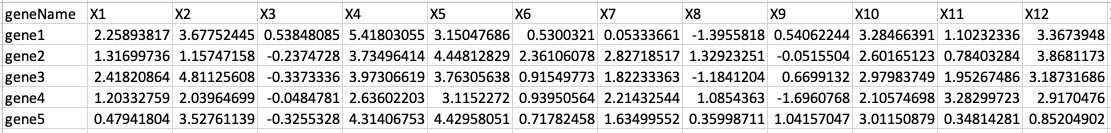

Figure 7: Sample expression data for circadian rhythmicity detection with repeated measurement

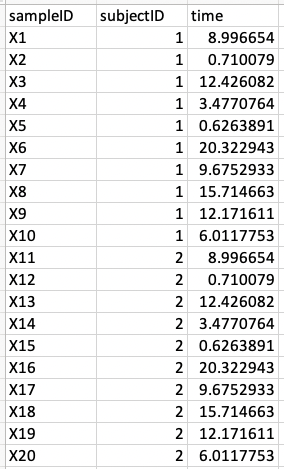

Figure 8: Sample phenotype data for circadian rhythmicity detection with repeated measurement

In the expression data, each row represents one gene, and each column represents one time point. The sample IDs (X1, X2, …) should match with the sample IDs in phenotype data. The unit of time variable in the phenotype data is hour(s). The subjectID in the phenotype data indicates subject ID for each sample ID.

**Input**

- Expression data: gene expression value data in .csv format (see Figure 6).
- Phenotype data: phenotype data in .csv format (see Figure 7).
- Period: period of gene expression value data, default is 24.
- Testing methods: available methods are Likelihood ratio test [1] and Cosinor2 [6]. Default is Likelihood ratio test.

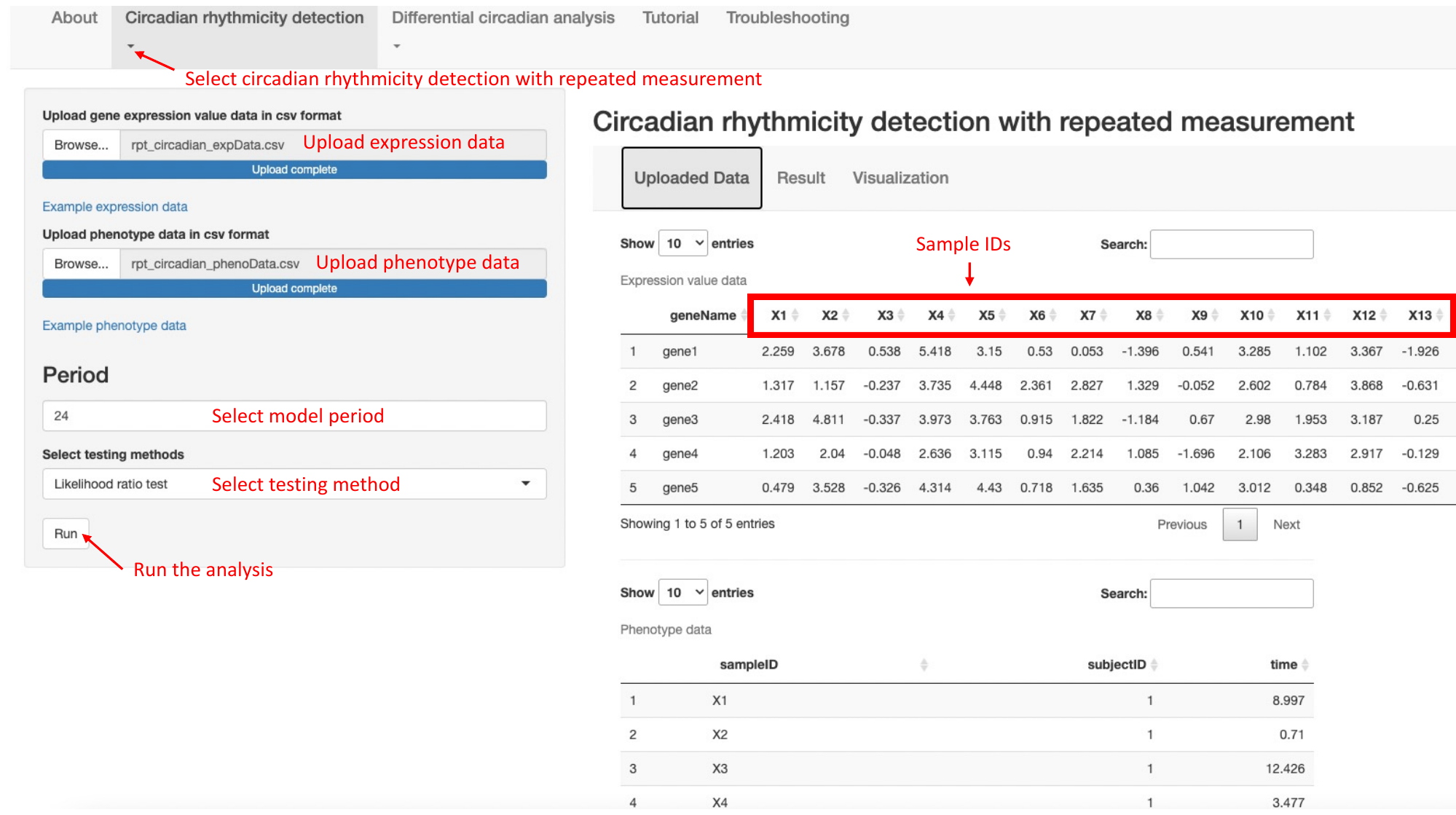

Figure 9: Circadian rhythmicity detection with repeated measurement after uploaded expression and phenotype data

**Run**

In left input panel, first upload both gene expression value data and phenotype data. Then input the period (default is 24) and select testing methods (see Figure 8). After running the analysis, click the ‘Result’ box in the right panel, it will show the estimates summary table. The ‘Download Result’ button allows you to down the result table (see Figure 9). In addition, click ‘Visualization’ box allows user to visualize the model fitting result. To plot the result, simply input interested gene name and select either overall model fitting plot or individual subject model fitting plot. Then click the ‘Plot’ button and download the plot with ‘Download Figure’ button (see Figure 10).

**Output**

A result table including cosine model parameter estimates and statistics (see Figure 9), detailed explanations are shown below:

- geneName: gene name.
- amplitude: estimated amplitude.
- peakTime: estimated peak time of the cosine curve.
- MESOR: estimated value of MESOR level (vertical shift).
- pvalue: p-value from the test.

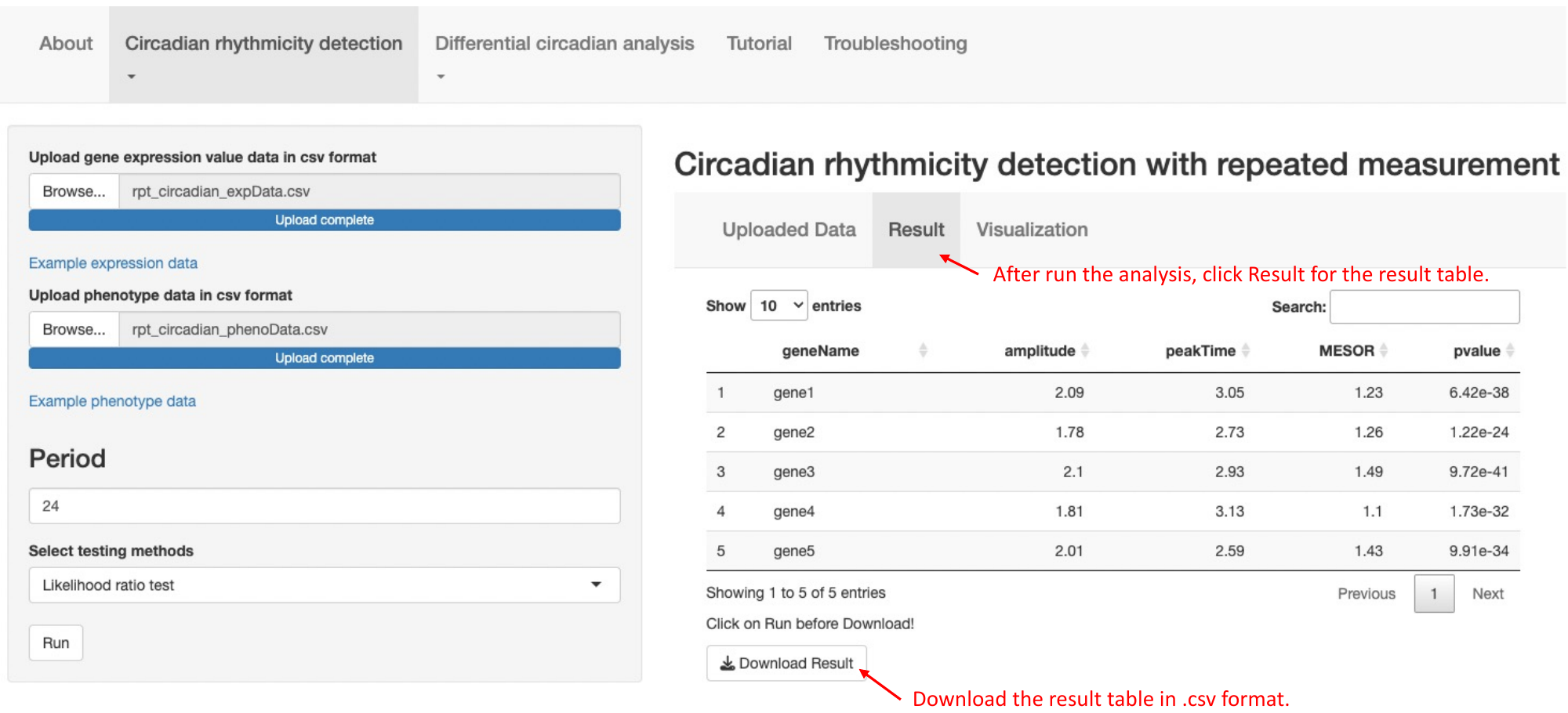

Figure 10: Result table of circadian rhythmicity detection with repeated measurement

Circadian gene plot: model fitting plot of interested circadian gene.

- Gene name: interested gene name.
- Plot type: either plot the fitted model for all subjects (‘overall’ option) or plot the models for each individual subject (‘individual’ option).

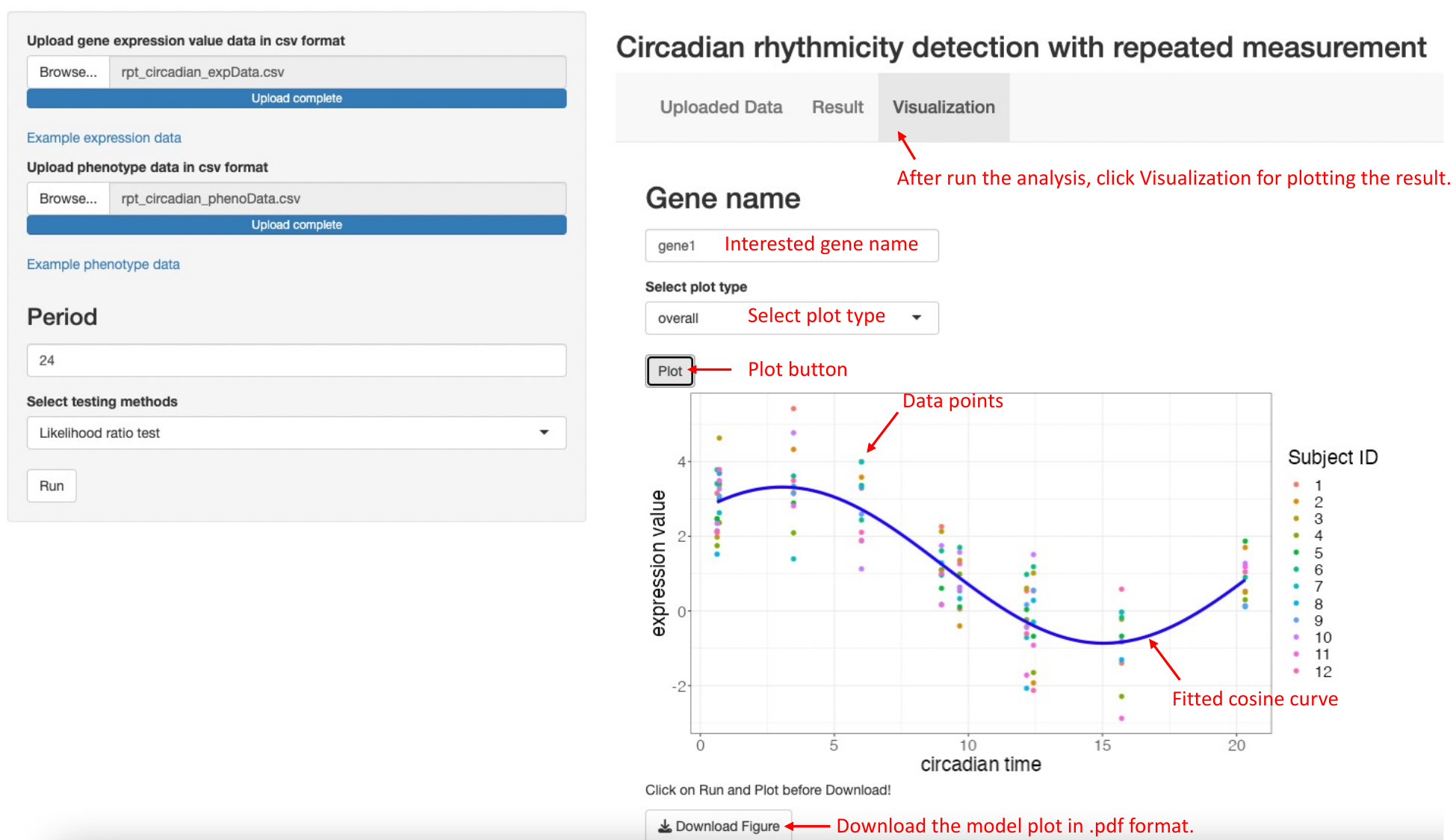

Figure 11: Visualization result of circadian rhythmicity detection with repeated measurement

#### Module 2: Differential circadian analysis

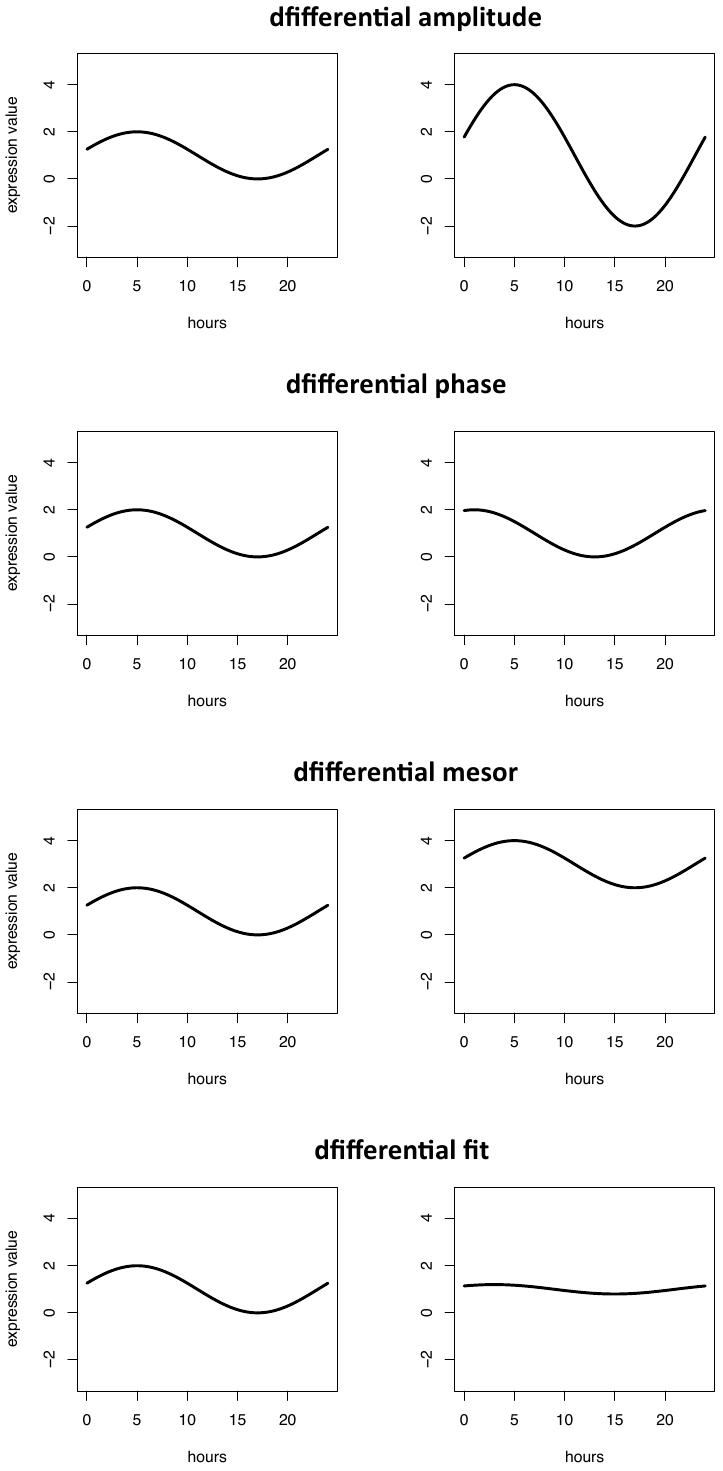

Figure 12: Four subtypes of differential circadian patterns

##### Figure 12 shows four subtypes of differential circadian patterns (differential amplitude, phase, MESOR and fit). For example, in terms of differential amplitude, the amplitudes between two conditions are different.

**Hypothesis testing**

Below are the null hypothesis and the alternative hypothesis for testing four categories of differential circadian patterns. For example, while testing differential amplitude, A_1_ is the amplitude for condition 1 and A_2_ is the amplitude for condition 2. A_c_ represent the common amplitude between condition 1 and condition 2 when under the null hypothesis that there is no differential amplitude.

1. Differential amplitude: $H_{0} : A_{1} = A_{2} = A_{c} v.s. H_{A} : A_{1}\neq A_{2}$.

2. Differential phase: $H_{0} : \varphi_{1} = \varphi_{2} = \varphi_{c} v.s. H_{A} : \varphi_{1}\neq\varphi_{2}$.

3. Differential basal level: $H_{0} : M_{1} = M_{2} = M_{c} v.s. H_{A} : M_{1}\neq M_{2}$.

4. Differential fit: $H_{0}:\sigma_{1}=\sigma_{2} v.s. H_{A}:\sigma_{1}\neq\sigma_{2}$.

Note that, as suggested by [7] Chen et al σ^2^ is used as a closely related quantity to quantify the goodness of fit.

##### a. with independent samples

**Data preparation**

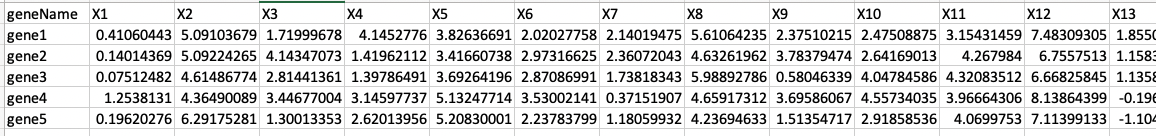

Figure 13: Sample expression data for differential circadian analysis with independent samples

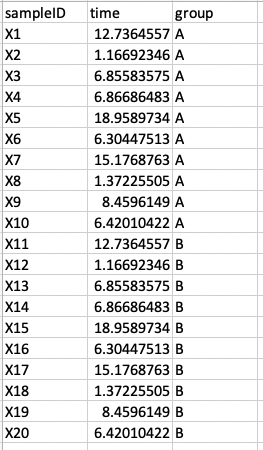

Figure 14: Sample phenotype data for differential circadian analysis with independent samples

In the expression data, each row represents one gene, and each column represents one time point. The sample IDs (X1, X2, …) should match with the sample IDs in phenotype data. The unit of time variable in phenotype data is hour(s). Group variable in the phenotype data indicates the group information for each sample.

**Input**

- Expression data: gene expression value data in .csv format (see Figure 11).
- Phenotype data: phenotype data in .csv format (see Figure 12).
- Period: period of gene expression value data, default is 24.
- Test types: type 1 is testing whether circadian rhythmicity is identical across two conditions; type 2 is testing whether subcategories of differential circadian patterns are different between two conditions.
- Test methods: selecting test method for the analysis. If type 1 is selected, available methods are likelihood ratio test [2] (default), HANOVA [3] and Limorhyde [4]. If type 2 is selected, available methods are likelihood ratio test (default) and CircaCompare [5].

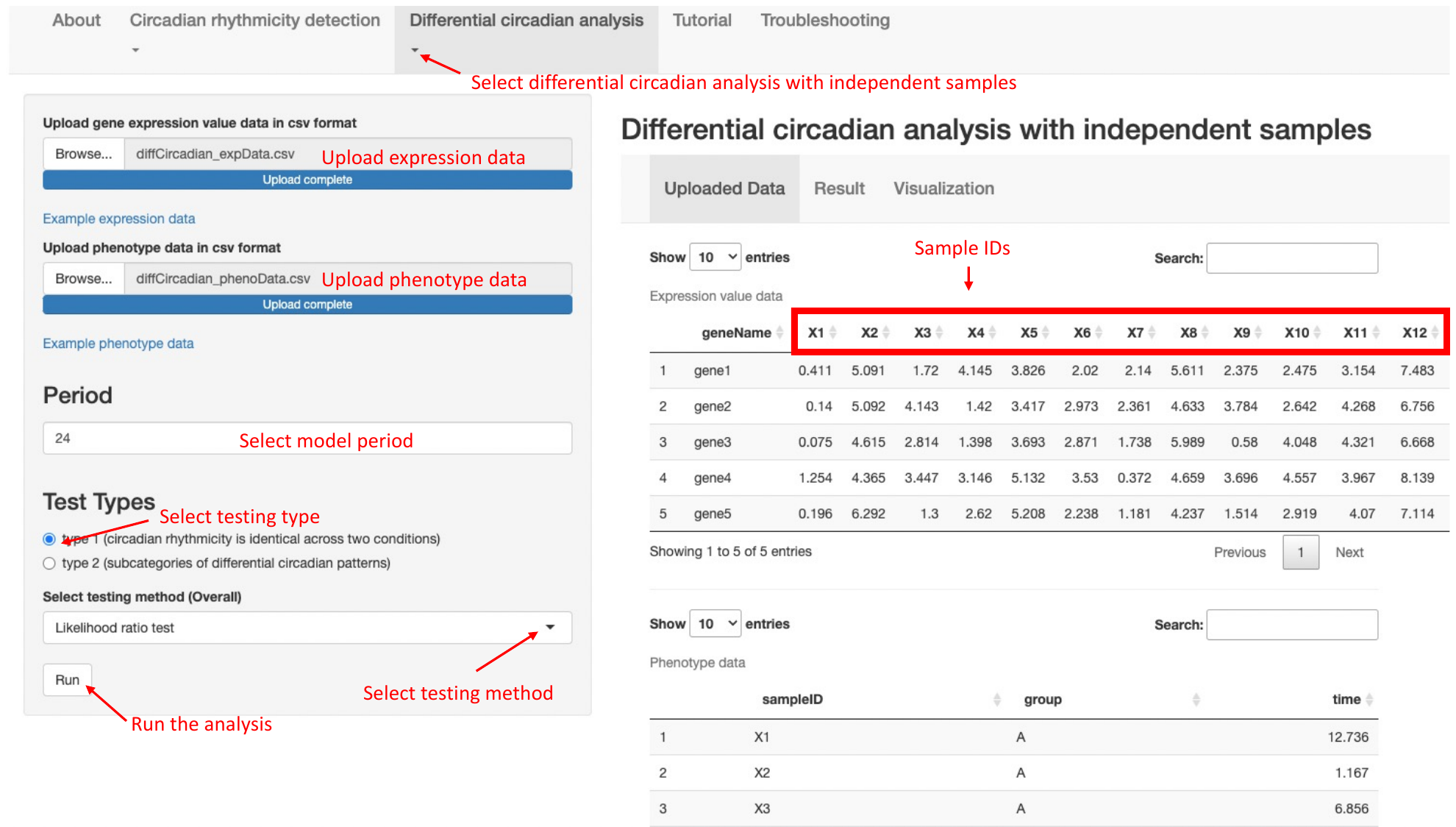

Figure 15: Differential circadian analysis with independent samples after uploaded expression and phenotype data

**Run**

In left input panel, first upload both gene expression value data and phenotype data. Then input the period (default is 24) and select test type and method (see Figure 13). After running the analysis, click the ‘Result’ box in the right panel, it will show the estimates summary table. The ‘Download Result’ button allows you to down the result table (see Figure 14). In addition, click ‘Visualization’ box allows user to visualize the model fitting result. To plot the result, simply input interested gene name then click the ‘Plot’ button and download the plot with ‘Download Figure’ button (see Figure 15).

**Output**

A result table including cosine model parameter estimates and statistics (see Figure 9: Result table of differential circadian analysis with independent samples), detailed explanations are shown below:

- geneName: gene name.
- amplitude_(group 1 name): estimated amplitude in group 1.
- amplitude_(group 2 name): estimated amplitude in group 2.
- peakTime_(group 1 name): estimated peak time of the cosine curve in group 1.
- peakTime_(group 2 name): estimated peak time of the cosine curve in group 2.
- MESOR_(group 1 name): estimated value of MESOR level (vertical shift) in group 1.
- MESOR_(group 2 name): estimated value of MESOR level (vertical shift) in group 2.
- pvalue: p-value from the (overall) test (Not available in if ‘Type 2’ is selected).

If Test Type is ‘type 2’, there will have following additional estimates:

- diff_amp_p: p-value from testing differential amplitude.
- diff_peak_p: p-value from testing differential peak time.
- diff_MESOR_p: p-value from testing differential MESOR.
- sigma2_(group 1 name): Variance estimate of group 1 data.
- sigma2_(group 2 name): Variance estimate of group 2 data.
- diff_fit_p: p-value from testing differential goodness of fitness.

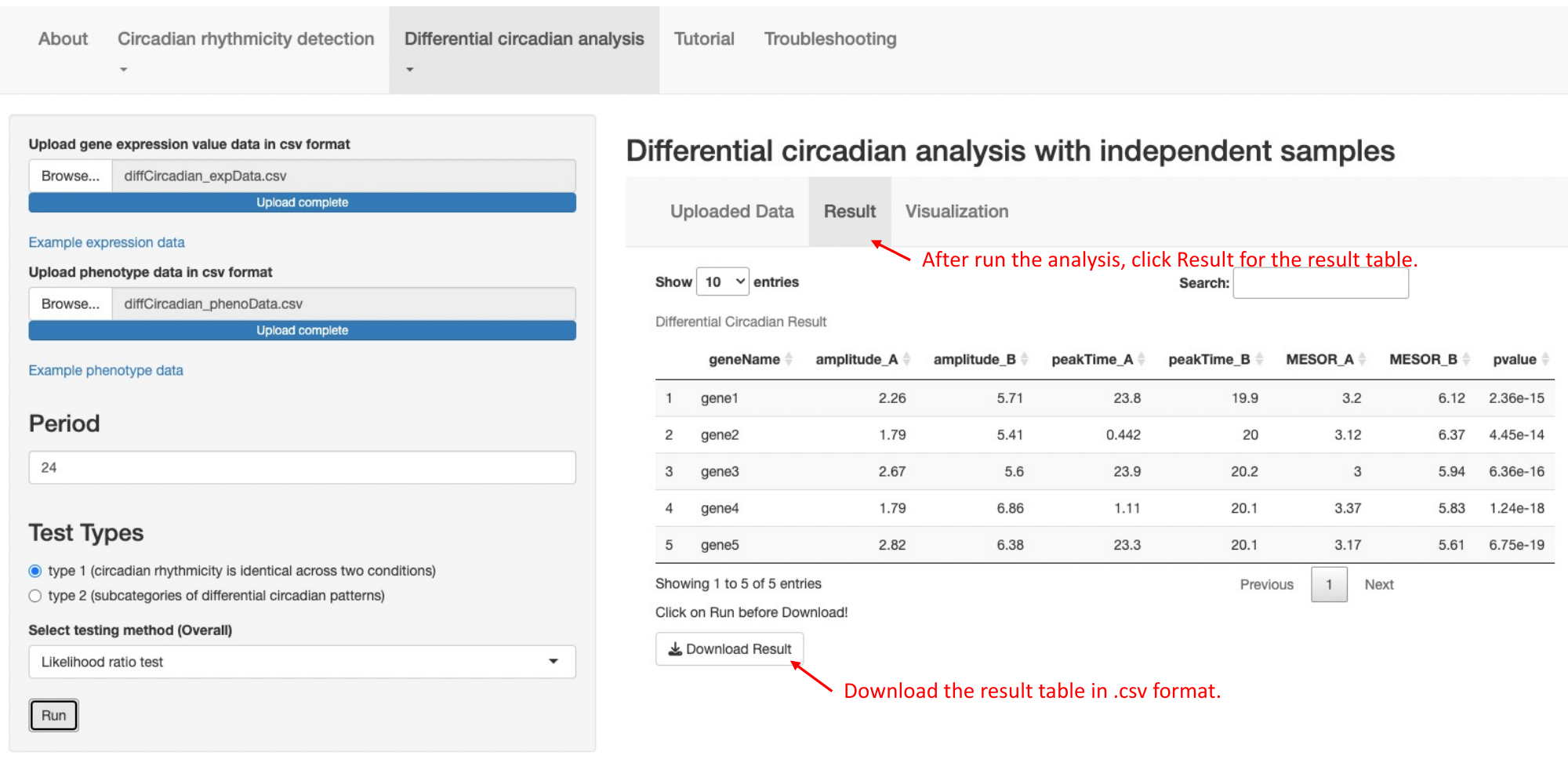

Figure 16: Result table of differential circadian analysis with independent samples

Differential circadian gene plot: model fitting plots by two experimental conditions with interested gene.

- Gene name: interested gene name.

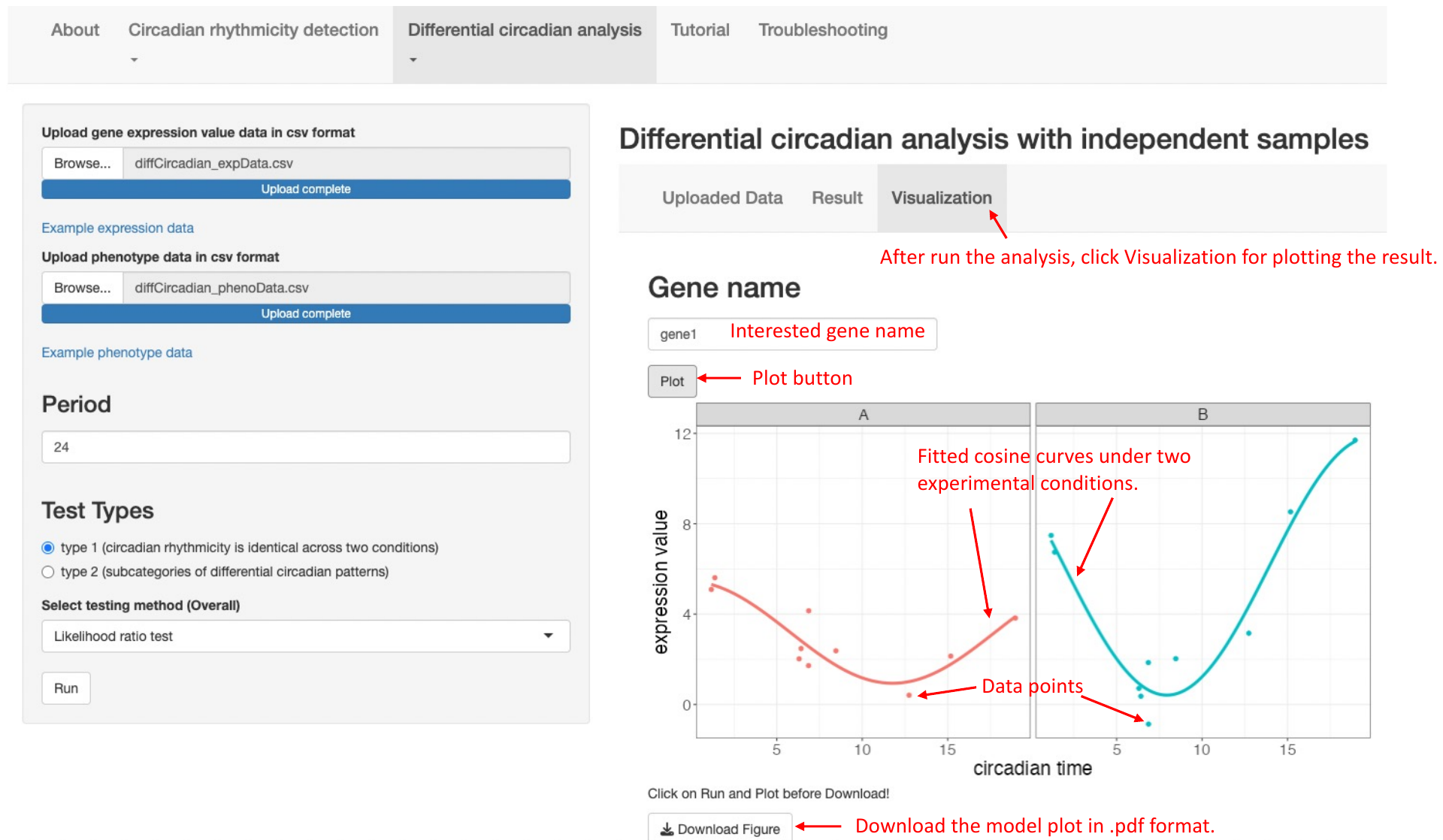

Figure 17: Visualization result of differential circadian analysis with independent samples

##### b. with repeated measurement

**Data preparation**

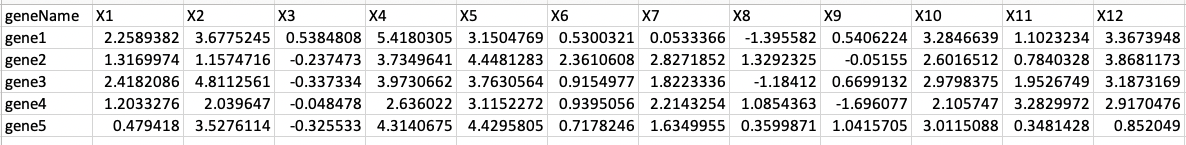

Figure 18: Sample expression data for differential circadian analysis with repeated measurement

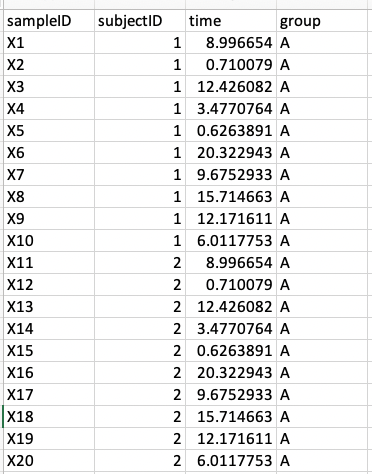

Figure 19: Sample phenotype data for differential circadian analysis with repeated measurement

In the expression data, each row represents one gene, and each column represents one time point. The sample IDs (X1, X2, …) should match with the sample IDs in phenotype data. The unit of time variable in phenotype data is hour(s). Group variable in the phenotype data indicates the group information for each sample. The subjectID in the phenotype data indicates subject ID for each sample ID.

**Input**

- Expression data: gene expression value data in .csv format (see Figure 16).
- Phenotype data: phenotype data in .csv format (see Figure 17).
- Period: period of gene expression value data, default is 24.
- Test types: type 1 is testing whether circadian rhythmicity is identical across two conditions; type 2 is testing whether subcategories of differential circadian patterns are different between two conditions.
- Test methods: selecting test method for the analysis. If type 1 is selected, the only available method is Likelihood ration test [2]. If type 2 is selected, the only available method is Cosinor2 [6].

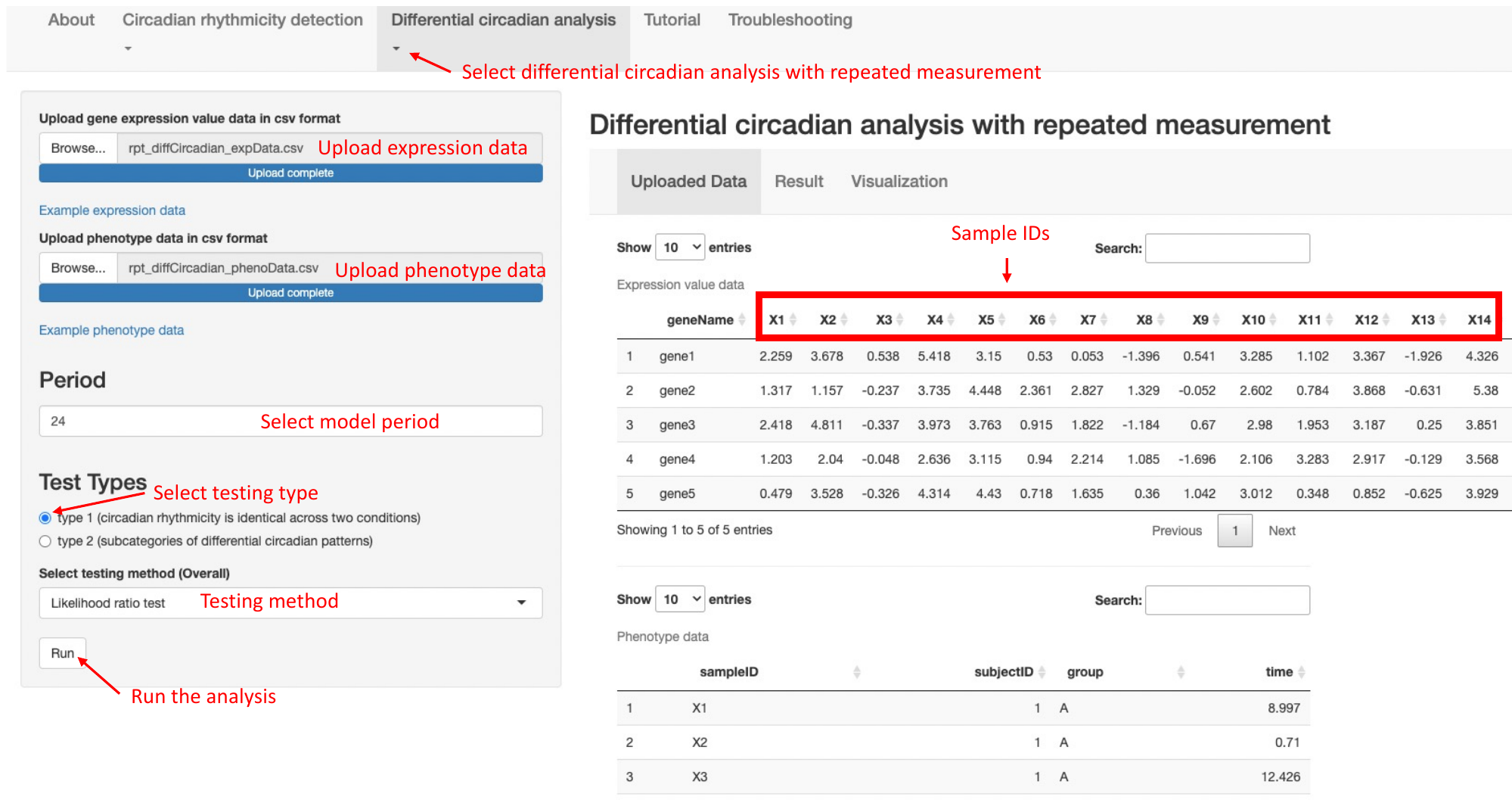

Figure 20: Differential circadian analysis with repeated measurement after uploaded expression and phenotype data

**Run**

In left input panel, first upload both gene expression value data and phenotype data. Then input the period (default is 24) and select test type and method (see Figure 18). After running the analysis, click the ‘Result’ box in the right panel, it will show the estimates summary table. The ‘Download Result’ button allows you to down the result table (see Figure 19). In addition, click ‘Visualization’ box allows user to visualize the model fitting result. To plot the result, simply input interested gene name and select either overall model fitting plot or individual subject model fitting plot. Then click the ‘Plot’ button and download the plot with ‘Download Figure’ button (see Figure 20).

**Output**

A result table including cosine model parameter estimates and statistics (see Figure 19), detailed explanations are shown below:

- geneName: gene name.
- amplitude_(group 1 name): estimated amplitude in group 1.
- amplitude_(group 2 name): estimated amplitude in group 2.
- peakTime_(group 1 name): estimated peak time of the cosine curve in group 1.
- peakTime_(group 2 name): estimated peak time of the cosine curve in group 2.
- MESOR_(group 1 name): estimated value of MESOR level (vertical shift) in group 1.
- MESOR_(group 2 name): estimated value of MESOR level (vertical shift) in group 2.
- pvalue: p-value from the (overall) test (Not available in if ‘Type 2’ is selected).

If Test Type is ‘type 2’, there will have following additional estimates:

- diff_amp_p: p-value from testing differential amplitude.
- diff_peak_p: p-value from testing differential peak time.
- diff_MESOR_p: p-value from testing differential MESOR.

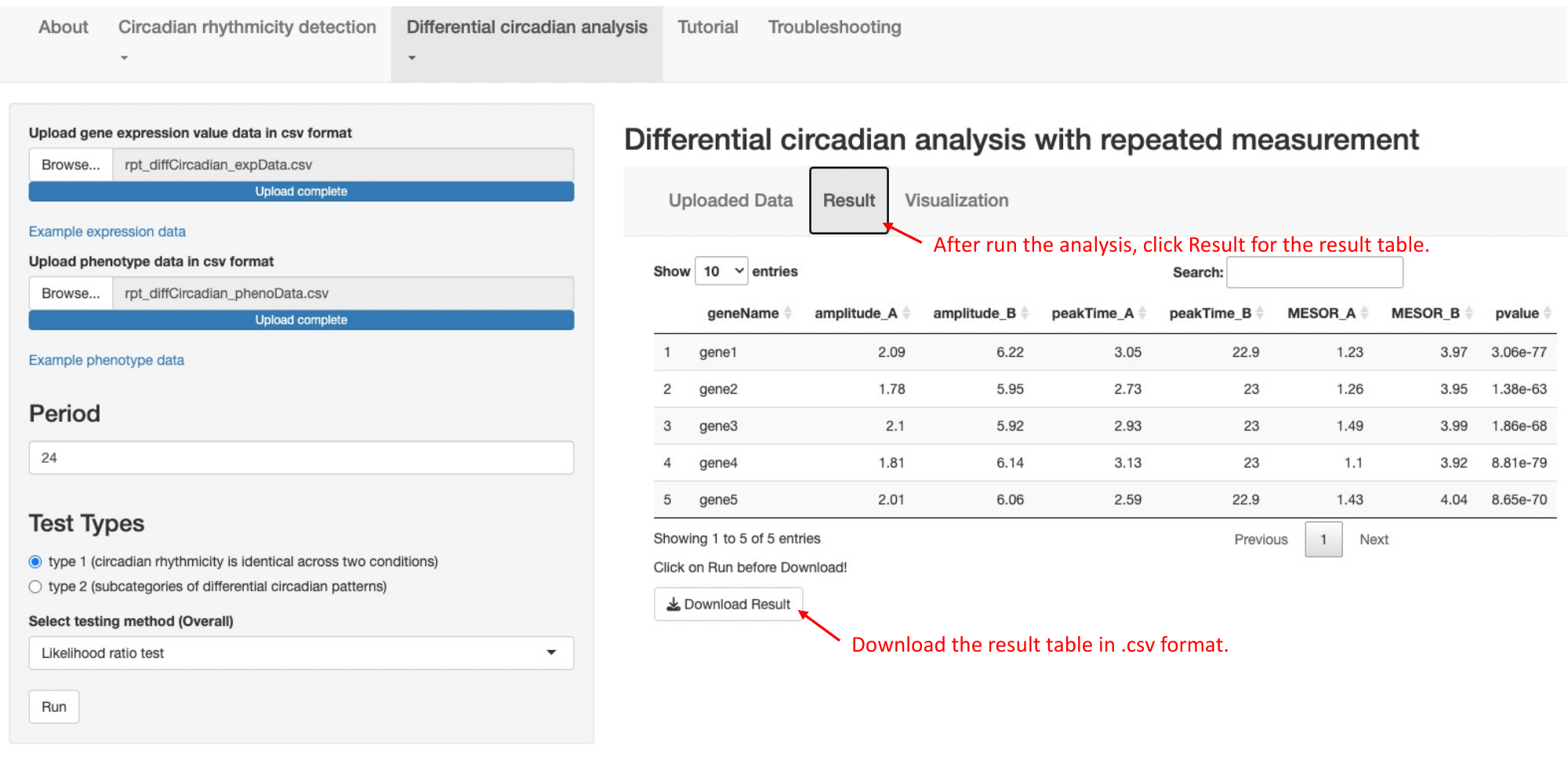

Figure 21: Result table of differential circadian analysis with repeated measurement

Differential circadian gene plot: model fitting plots by two experimental conditions with interested gene.

- Gene name: interested gene name.
- Plot type: either plot the fitted model for all subjects (‘overall’ option) or plot the models for each individual subject (‘individual’ option).

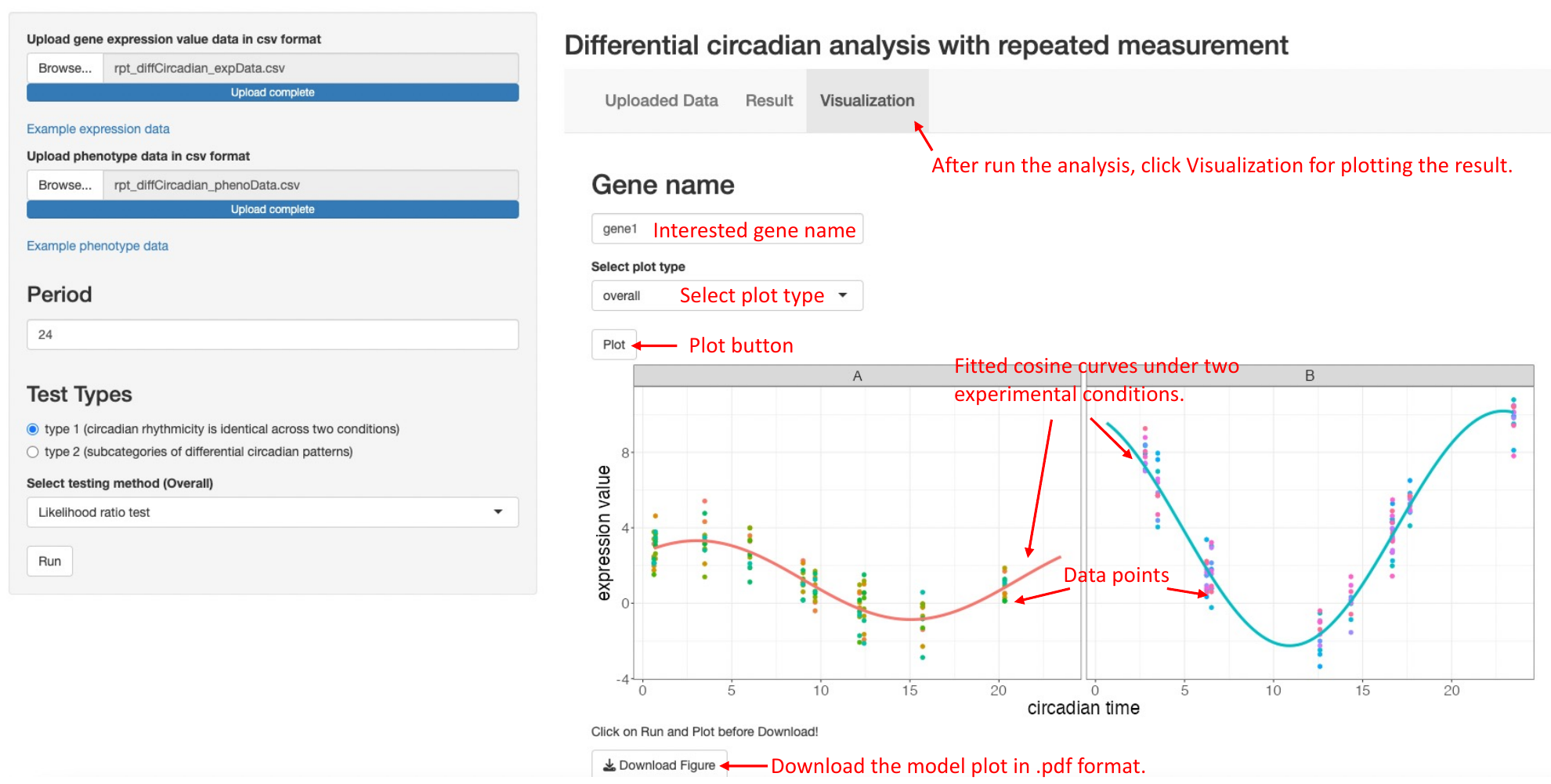

Figure 22: Visualization result of differential circadian analysis with repeated measurement

### 4. Reference

###### **Likelihood ratio test**

[1] Ding, H., Meng, L., Liu, A. C., Gumz, M. L., Bryant, A. J., Mcclung, C. A., Tseng, G. C., Esser, K. A., and Huo, Z. (2021). Likelihood-based tests for detecting circadian rhythmicity and differential circadian patterns in transcriptomic applications. Briefings in Bioinformatics, 22(6). bbab224

[2] Ding, H., Meng, L., Xing, C., Esser, K. A., & Huo, Z. (2022). Statistical Methods for Detecting Circadian Rhythmicity and Differential Circadian Patterns with Repeated Measurement in Transcriptomic Applications. bioRxiv, 2022.06. 05.494875

###### **HANOVA**

[3] Thaben, P. F. and Westermark, P. O. (2016). Differential rhythmicity: detecting altered rhythmicity in biological data. Bioinformatics, 32(18):2800–2808

###### **Limorhyde**

[4] Singer, J. M. and Hughey, J. J. (2019). Limorhyde: a flexible approach for differential analysis of rhythmic transcriptome data. Journal of biological rhythms, 34(1):5–18

###### **CircaCompare**

[5] Parsons, R., Parsons, R., Garner, N., Oster, H., and Rawashdeh, O. (2020). Circacompare: a method to estimate and statistically support differences in mesor, amplitude and phase, between circadian rhythms. Bioinformatics, 36(4):1208–1212

###### **Cosinor2**

###### [6] Cornelissen, G. (2014). Cosinor-based rhythmometry. Theoretical Biology and Medical Modelling, 11(1):16

**Others**

[7] Chen, C.-Y., Logan, R. W., Ma, T., Lewis, D. A., Tseng, G. C., Sibille, E., and McClung, C. A. (2016). Effects of aging on circadian patterns of gene expression in the human prefrontal cortex. Proceedings of the National Academy of Sciences, 113(1):206–211
